# *destiny* – diffusion maps for large-scale single-cell data in R

**DOI:** 10.1101/023309

**Authors:** Philipp Angerer, Laleh Haghverdi, Maren Büttner, Fabian J. Theis, Carsten Marr, Florian Buettner

## Abstract

**Summary:** Diffusion maps are a spectral method for non-linear dimension reduction and have recently been adapted for the visualization of single cell expression data. Here we present *destiny*, an efficient R implementation of the diffusion map algorithm. Our package includes a single-cell specific noise model allowing for missing and censored values. In contrast to previous implementations, we further present an efficient nearest-neighbour approximation that allows for the processing of hundreds of thousands of cells and a functionality for projecting new data on existing diffusion maps. We exemplarily apply *destiny* to a recent time-resolved mass cytometry dataset of cellular reprogramming.

**Availability and implementation:** *destiny* is an open-source R/Bioconductor package http://bioconductor.org/packages/ destiny also available at https://www.helmholtz-muenchen.de/icb/destiny. A detailed vignette describing functions and workflows is provided with the package.

## 1 INTRODUCTION

Recent technological advances allow for the profiling of individual cells, using methods such as single-cell RNA-seq, single-cell RT qPCR or cyTOF (Vargas Roditi and Claassen, 2015) These techniques have been used successfully to study stem cell differentiation with time-resolved single-cell experiments, where individual cells are collected at different absolute times within the differentiation process and profiled. While differentiation is a smooth but nonlinear process (Buettner and Theis, 2012; Haghverdi, Buettner, and Theis, 2015) involving continuous changes of the overall transcriptional state, standard methods for visualizing such data are either based on linear methods such as Principal Component Analysis and Independent Components Analysis or they use clustering techniques not accounting for the smooth nature of the data.

In contrast, diffusion maps – initially designed by Coifman, Lafon, et al. (2005) for dimensionality reduction in image processing – recover a distance measure between each pair of data points (cells) in a low dimensional space that is based on the transition probability from one cell to the other through several paths of a random walk. Diffusion maps are especially suited for analyzing single-cell gene expression data from differentiation experiments (such as time-course experiments) for three reasons. First, they preserve the global relations between data points. This feature makes it possible to reconstruct developmental traces by re-ordering the asynchronously differentiating cells according to their internal differentiation state. Second, the notion of diffusion distance is robust to noise, which is ubiquitous in single-cell data. Third, by normalizing for sampling density, diffusion maps become insensitive to the distribution of the data points (i. e. sampling density), which aids the detection of rare cell populations.

Here, we present a user friendly R implementation of diffusion maps including previously proposed adaptations to single cell data (Haghverdi, Buettner, and Theis, 2015) as well as novel functionality. The latter includes approximations allowing for the visualisation of large data sets and the projection of new data on existing maps.

## 2 DESCRIPTION: THE *DESTINY* PACKAGE

### 2.1 Algorithm

As input, *destiny* accepts an expression matrix or data structure extended with annotation columns. Gene expression data should be pre-processed and normalized using standard workflows (see Supplementary text S1) before generating the diffusion map. *destiny* calculates cell-to-cell transition probabilities based on a Gaussian kernel with width *σ* to create a sparse transition probability matrix *M*. If the user does not specify *σ, destiny* employs an estimation heuristic to derive this parameter (see Supplementary Text S2). In contrast to other implementations, *destiny* allows for the visualisation of hundreds of thousands of cells by only using distances to the *k* nearest neighbors of each cell for the estimation of *M* (see Supplementary Text S2). Optionally *destiny* uses an application-specific noise model for censored and missing values in the dataset (see Figure S1). An eigendecomposition is performed on *M* after density normalization, considering only transition probabilities between different cells. By rotating *M*, a symmetric adjoint matrix can be used for a faster and more robust eigendecomposition (Coifman, Kevrekidis, et al., 2008) The resulting data-structure contains the eigenvectors with decreasing eigenvalues as numbered diffusion components, the input parameters and a reference to the data.

### 2.2 Visualization and projection of new data

This data-structure can be easily plotted and colored using the parameters of provided plot methods. An automatic color legend integrated into R’s palette system facilitates the generation of publication-quality plots. A further new feature in *destiny* is the ability to integrate new experimental data in an already computed diffusion map. *destiny* provides a projection function to generate the coordinates for the new data without recalculating the diffusion map by computing the transition probabilities from new data points to the existing data points (see Supplementary Text S3).

## 3 APPLICATION

We applied *destiny* to four single-cell datasets of different size (hundreds to hundreds of thousands of cells) and characteristics (qRT-PCR, RNA-Seq and mass cytometry, see Supplementary Table S1). We first estimate the optimal *σ* that matches the intrinsic dimensionality of the data (Fig. 1A and Supplementary Figs. S2A and S3A). Using a scree plot (Fig. 1B and Supplementary Figs. S2B, S3B, and S4A), the relevant diffusion components can be identified. However, for big datasets as the mass cytometry data from Zunder et al. (2015) with 256,000 cells and 36 markers, corresponding Eigenvalues decrease smoothly. Although only a part of the intrinsic dimensionality can be represented in a 3D plot, the diffustion map reveals interesting properties of the reprogramming dynamics (Fig. 1C and Supplementary Fig. S5). We compared *destiny’s* performance to other implementations, including our own in MATLAB (based on Maggioni code^1^, published with Haghverdi, Buettner, and Theis, 2015) and the diffusionMap R package (Richards, 2014) *destiny* performs similarly well for small datasets, while outperforming other implementations for large datasets (see Supplementary Table S1).

**Figure 1.**
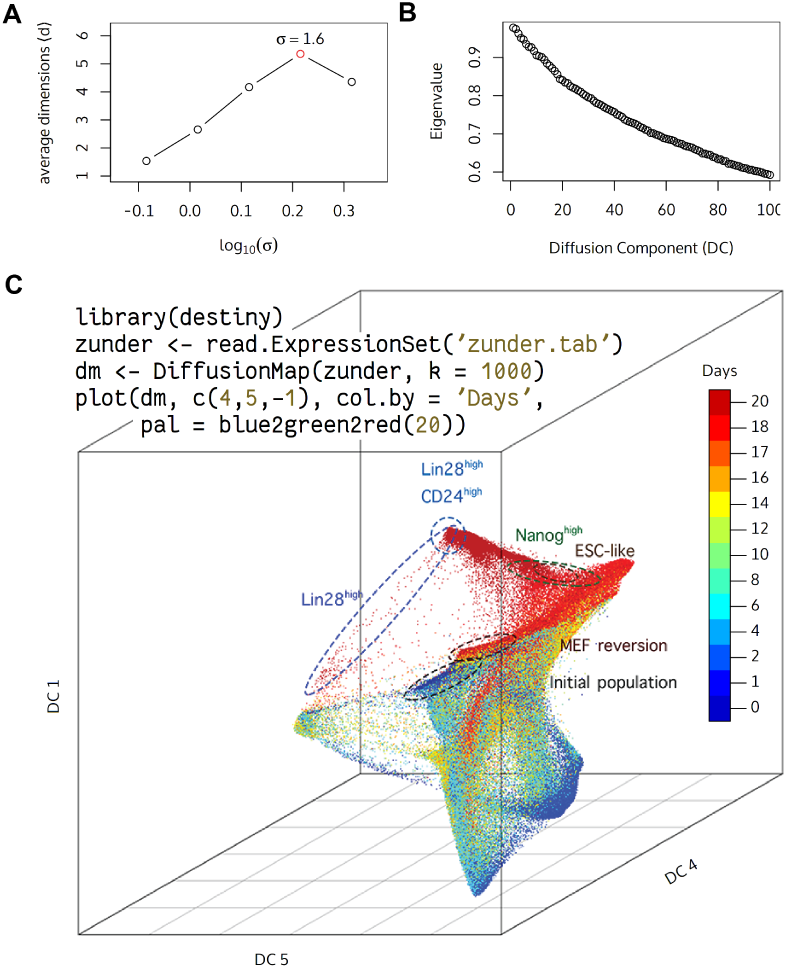
*destiny* applied to the mass cytometry reprogramming dataset of Zunder et al. (2015) with 36 markers and 256,000 cells. A) The optimal Gaussian kernel width *σ*. B) The Eigenvalues of the first 100 diffusion components decrease smoothly, indicating a large intrinsic dimensionality of the data. C) The initial population of mouse embryonic fibroblasts (MEFs, blue) is reprogrammed and profiled over 20 days. While a final cell population expressing stem cell markers is clearly separated, cells that revert to the MEF state are found proximal to the initial population in the diffusion map. (inset:) *destiny* code to generate the diffusion map

## 4 DISCUSSION AND CONCLUSION

We present a user-friendly R package of the diffusion map algorithm adapted to single-cell gene expression data and include new features for efficient handling of large datasets and a projection functionality for new data. We illustrate the capabilities of our package by visualizing gene expression data of 250,000 cells and show that our package is able to reveal continuous state transitions. Together with an easy to use interface this facilitates the application of diffusion map as new analysis tool for single-cell gene expression data.

## ACKNOWLEDGEMENT

We thank Chris McGinnis (Seattle, USA) and Vicki Moignard (Cambridge, UK) for helpful comments on *destiny.*

*Funding:* Supported by the UK Medical Research Council (Career Development Award to FB) and the ERC (starting grant Latent-Causes to FJT). MB is supported by a DFG Fellowship through the Graduate School of Quantitative Biosciences Munich (QBM).

## Supplementary Figures and Text for *“destiny* – diffusion maps for single-cell time-course data”

### SUPPLEMENTARY TEXT

#### S1 Preprocessing

The data properties required by destiny include a variance not spanning many orders of magnitude due to the global *σ*. Therefore, RNAseq count data should be adapted using a variance-stabilizing transformation like a logarithm or the square root (Stegle, Teichmann, and Marioni, 2015) For single-cell qPCR data, expression values can be normalized by dividing (or subtracting in the case of logarithmic values) by housekeeper gene expression levels. However, as the expression of housekeeping genes is also stochastic, it is not clear whether such normalization is beneficial. Consequently, if such housekeeping normalization is performed, it is crucial to take the mean of several genes.

#### S2 Parameter selection

For the choice of the Gaussian kernel width, we pick a *σ* close to the intrinsic dimensionality of data. Such choice of the kernel width ensures a maximum connectivity of the data point as a graph while restricting the use of euclidean distances in the Gaussian kernel to local proximities where such distances are valid. How the intrinsic dimensionality of data can be approximated has been shown in Haghverdi, Buettner, and Theis (2015)

If the nearest neighbors approximation is used, the input parameter *k* controls the number of nearest neighbours for each cell to be considered. Guideline for *k* is a small enough number to make the computation cost limited, but not too small to alter the connectivity of data as a graph, which would result in a noisy embedding. A typical *k* is between 200 and 1000 cells.

#### S3 Projection of new data

Projection of new data onto existing diffusion components is done by first computing the transition probability matrix *M*′ from new data points to the existing data points. The same transformation that brought the existing data points’ transition matrix M to the low dimensional embedding, *M* × *C*, is applied to the newly built (*n_new_* × *n_init_*) transition matrix *M*′, *M*′ × *C*.

This way we provide an approximation for the diffusion distances of new points to all initial points. This approach is however inefficient if newly introduced points are too distant to any part of the initially existing population, since in a better approximation the new points will alter the position of initial points on the low dimensional map as well (Homrighausen and McDonald, 2011)

### SUPPLEMENTARY FIGURES AND TABLES

All scree plots (S2B, S3B, and S4A) correspond to a Markov chain length t = 1.

**Table S1.** Runtime comparison of *destiny* with previously available diffusion map implementations for three different single cell data sets. *destiny* utilizes sanity checking and imputing, slightly increasing runtime with respect to a fast MATLAB implementation. The fast MATLAB version does not include the censoring and missing value models. The data from Guo et al. (2010) Moignard et al. (2015) and Trapnell et al. (2014) have been processed on a 2GHz laptop with 8GB memory. As explained in S4, the latter was created using 20 singular values (SVs) of the data. The data from Zunder et al. (2015) have been processed on a 1.4GHz server with 954GB memory. The measurements for diffusionMap are always done with full distance matrices due to a lack of nearest neighbor approximation.

| | Guo et al. (2010)<br>(429 cells × 48 genes) | Guo et al. (2010)<br>(429 cells × 48 genes)<br>accounting for censored<br>& missing values | Moignard et al. (2015)<br>(3934 cells × 46 genes) | Trapnell et al. (2014)<br>(271 cells × 20 SVs) | Zunder et al. (2015)<br>(256k cells × 36 marker) |
| --- | --- | --- | --- | --- | --- |
| MATLAB fast | 0.3s | NA | 4.3s | 0.6s | >24h |
| MATLAB | 21s | 94s | >1h | 4.7s | NA (out of memory) |
| R *destiny* | 0.5s | 1.4s | 6.0s ($k=500$) | 0.2s | 1.4h ($k=1000$) |
| R diffusionMap | 0.4s | NA | 21s | 0.2s | NA (out of memory) |

**Figure S1.**
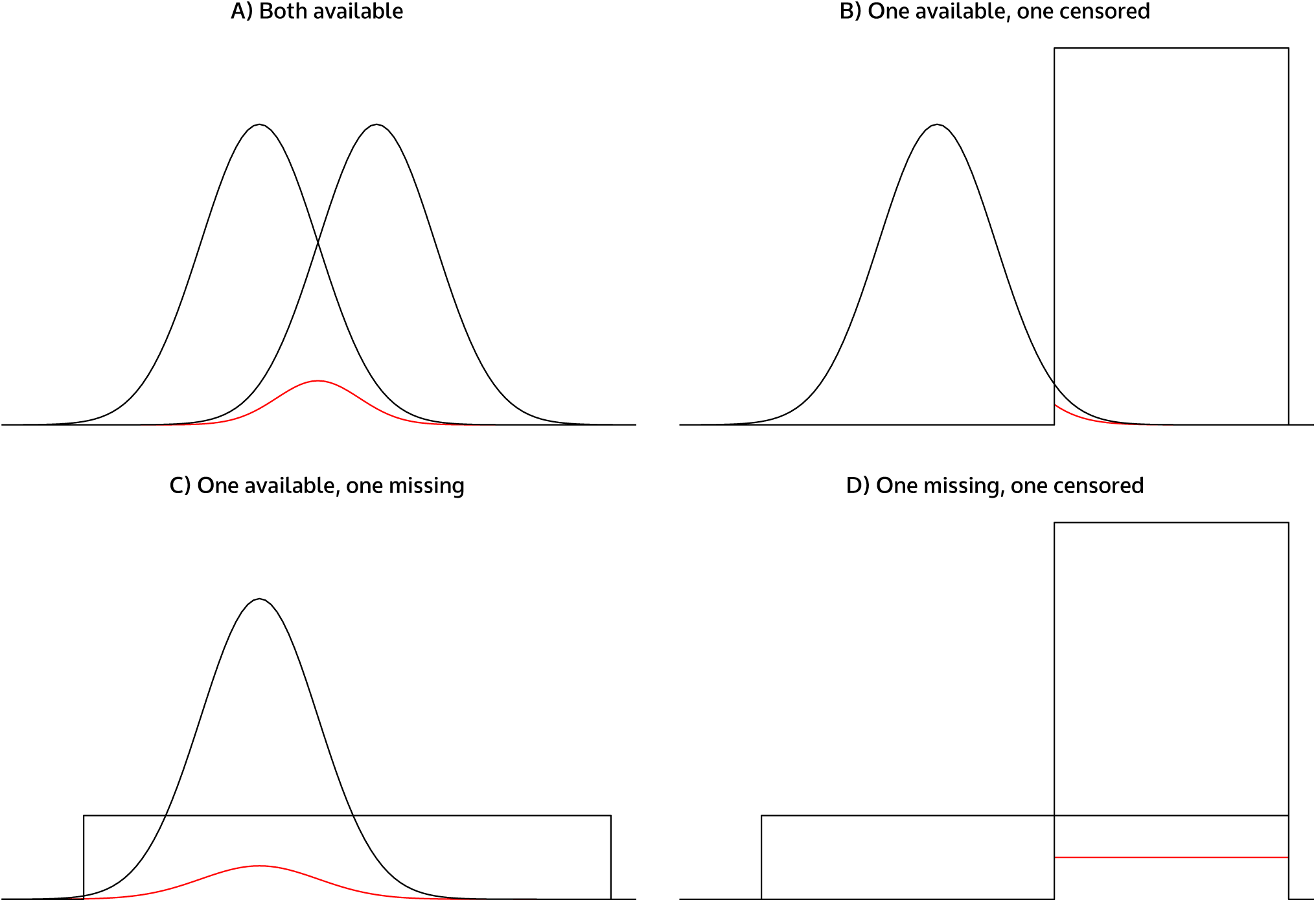
Transition probability models used for censored and missing values. A) The measurement of the current gene product is available for both cells. B) One measurement is missing. C) One measurement is set to the level of detection (i. e. censored) D) One measurement is missing, the other one censored.

**Figure S2.**
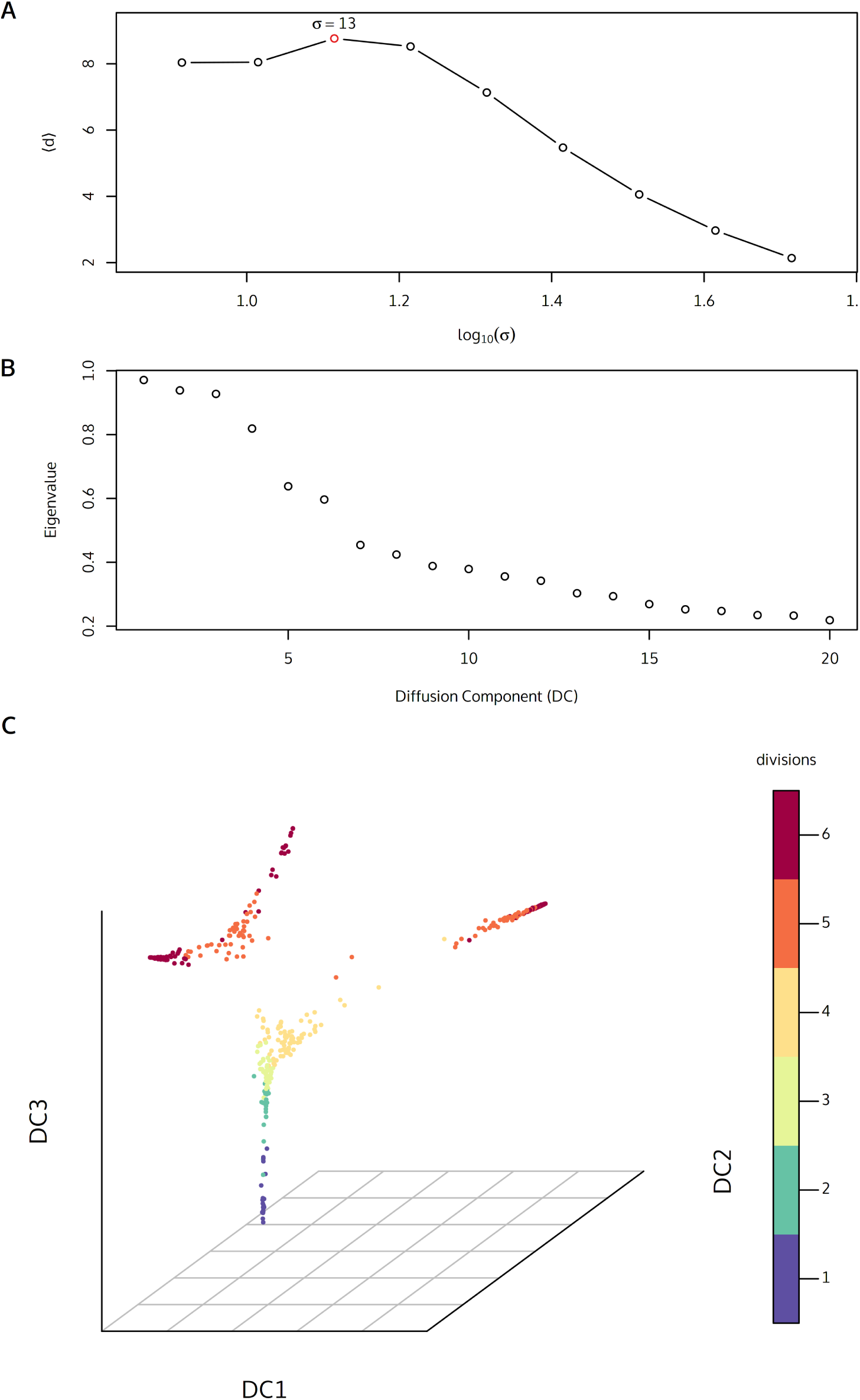
Diffusion map on single-cell qRT-PCR data of mouse embryonic cells from oocyte to 64-cell stage (Guo et al., 2010) A) Determination of the optimal Gaussian kernel width *σ*. B)The Eigenvalues of the first twenty diffusion components. The gap after the first three eigenvalues indicates an adequate dimension reduction on the first diffusion components. C) The *destiny* output for 48 genes from 429 cells arranges the cells in branches. Two different branching events can be identified leading to three clearly separated cell types in the 64-cell stage (6 divisions).

**Figure S3.**
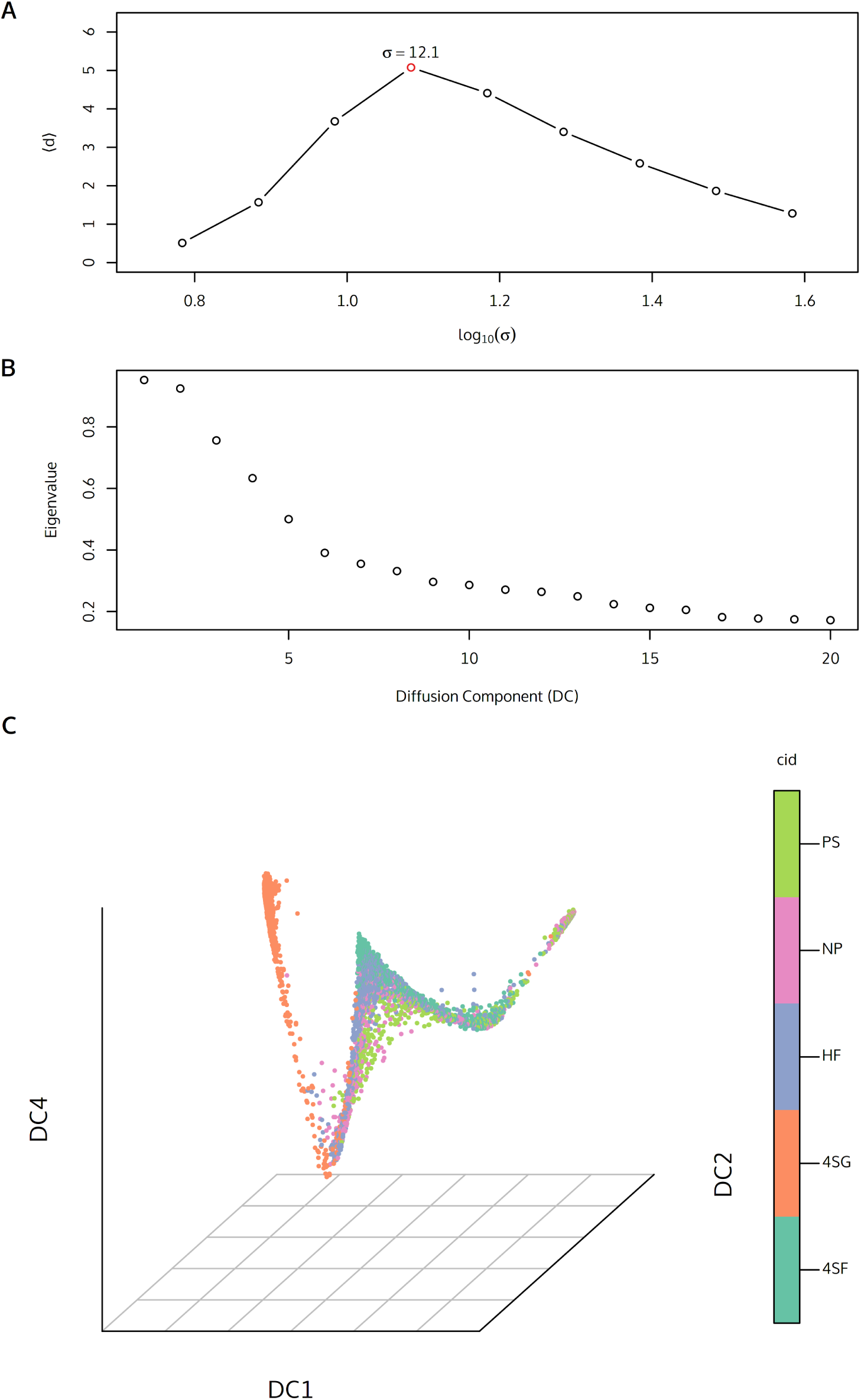
Diffusion map on single-cell qRT-PCR data of mouse embryonic cells with blood-forming potential (Moignard et al., 2015) A) Determination of the optimal Gaussian kernel width *σ*. B) The Eigenvalues of the first twenty diffusion components. The decrease of the Eigenvalues after the third component indicates C) The *destiny* output for 46 genes from 3934 cells. The diffusion trajectories *destiny* created correspond to the developmental stages of the progenitor cells (see Moignard et al. (2015)).

**Figure S4.**
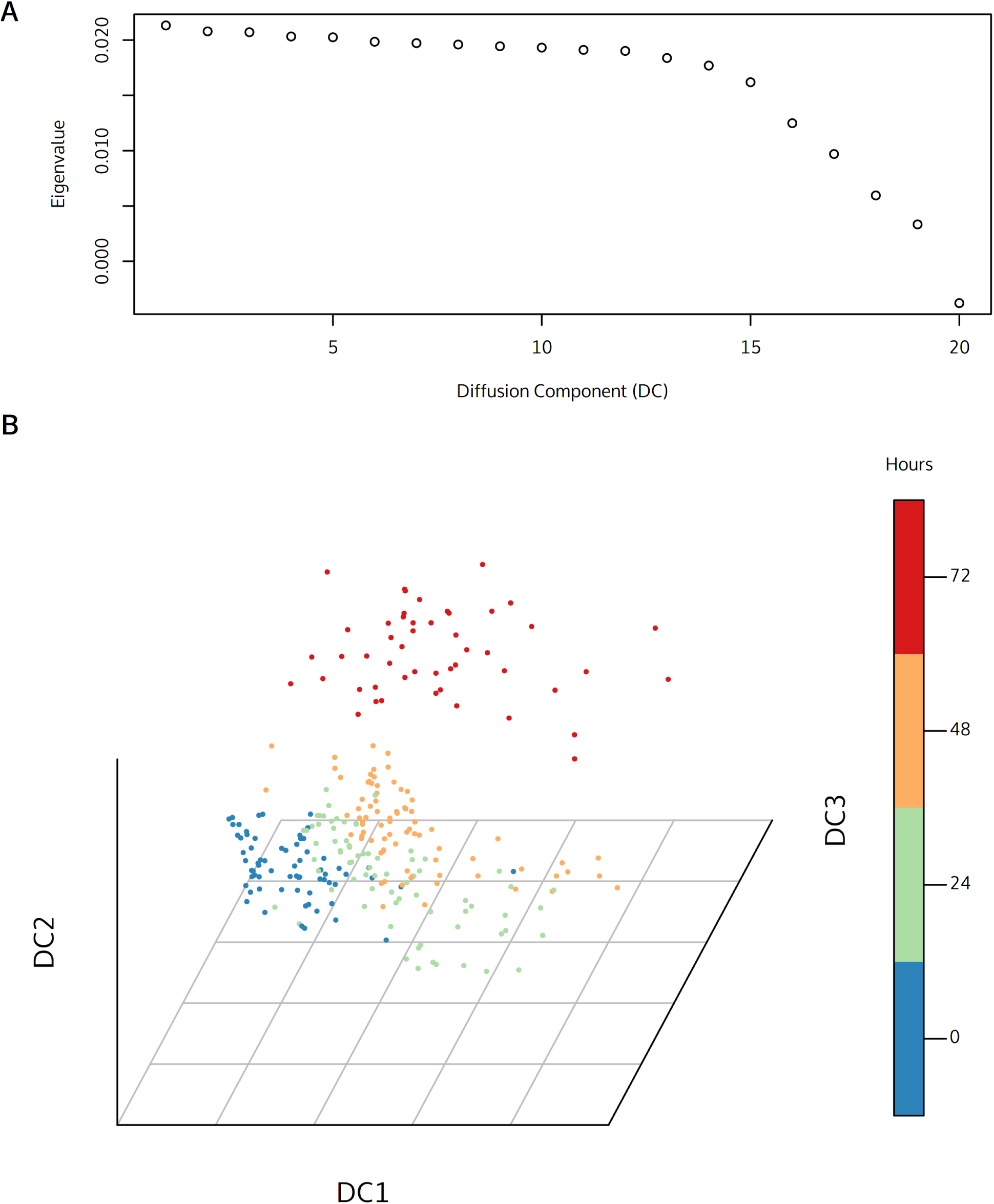
Diffusion map of the human skeletal muscle myoblasts RNA-Seq dataset from Trapnell et al. (2014) A) The Eigenvalues of the first twenty diffusion components B) The diffusion map computed from the first 20 singular values (SVs) of the RNA-Seq counts for 271 cells and 47192 genes. The sigma parameter was manually specified to 0.39.

**Figure S5.**
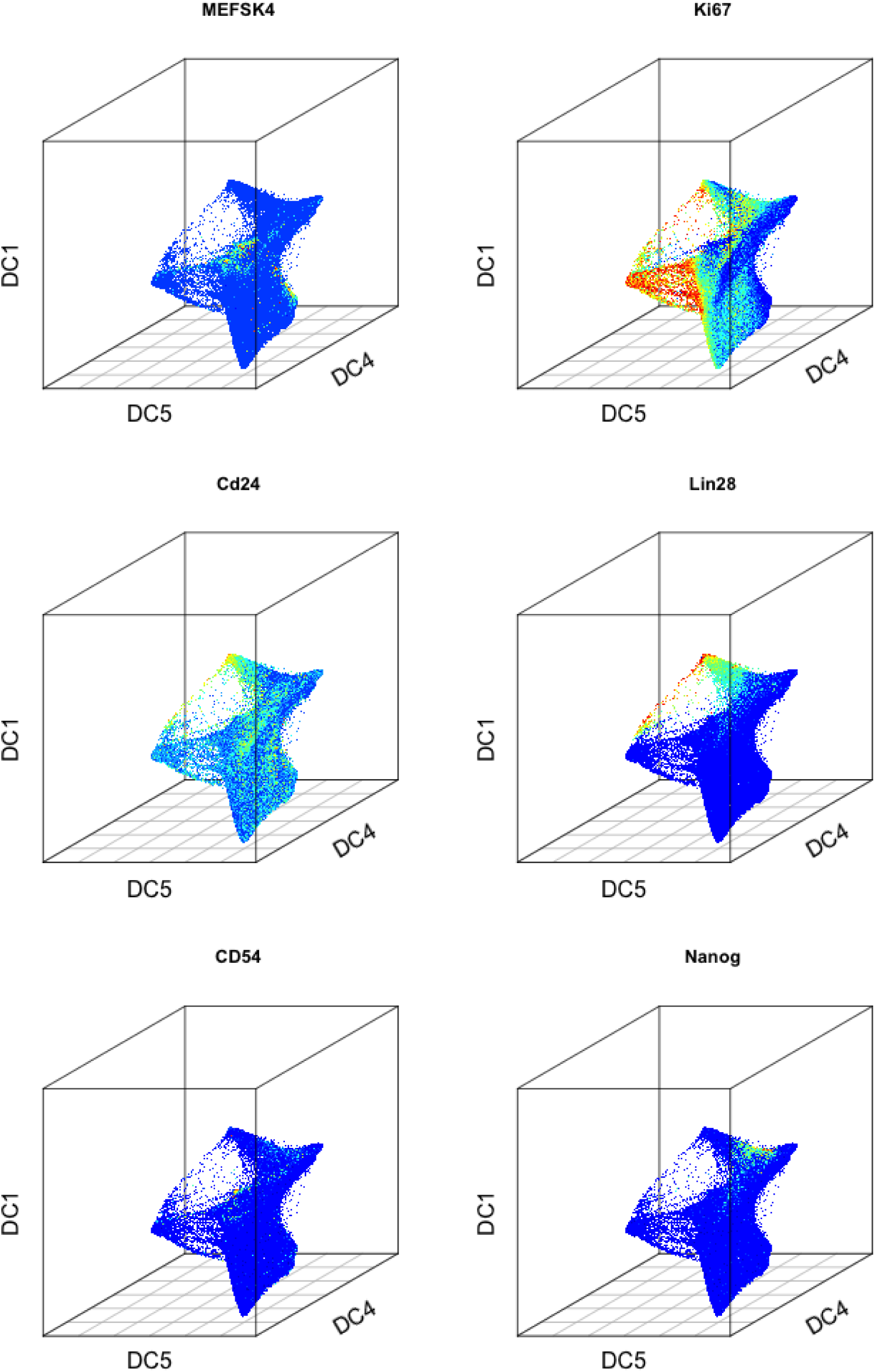
Expression of 6 intracellular and surface markers mapped on the diffusion map of the reprogramming dataset from Zunder et al. (2015) (see Figure 1). Coloring normalized on the log-scale (low - dark blue, medium - green, high - red).

## Footnotes

1 http://www.math.duke.edu/~mauro/code.html

